# Shifted balance of dorsal versus ventral striatal communication with frontal reward and regulatory regions in cannabis dependence

**DOI:** 10.1101/282939

**Authors:** Zhou Feng, Zimmermann Kaeli, Xin Fei, Dirk Scheele, Wolfgang Dau, Markus Banger, Bernd Weber, René Hurlemann, Keith M Kendrick, Benjamin Becker

## Abstract

The transition from voluntary to addictive behavior is characterized by a loss of regulatory control in favor of reward driven behavior. Animal models indicate that this process is neurally underpinned by a shift in ventral to dorsal striatal control of behavior, however this shift has not been directly examined in humans. Against this background the present resting state fMRI study employed a two-step approach to (1) precisely map striatal alterations using a novel, data-driven network classification strategy combining Intrinsic Connectivity Contrast (ICC) with Multivoxel Pattern Analysis (MVPA) and, (2) to determine whether a ventral to dorsal striatal shift in connectivity with reward and and regulatory control regions can be observed in abstinent (28 days) male cannabis-dependent individuals (n = 24) relative to matched controls (n = 28). Network classification revealed that the groups can be reliably discriminated by global connectivity profiles of two striatal regions that mapped onto the ventral (nucleus accumbens) and dorsal striatum (caudate). Subsequent functional connectivity analysis demonstrated a relative shift between ventral and dorsal striatal communication with fronto-limbic regions that have been consistently involved in reward processing (rostral ACC) and executive / regulatory functions (dorsomedial PFC). Specifically, in the cannabis dependent subjects connectivity between the ventral striatum with the rostral ACC increased, whereas both striatal regions were uncoupled from the regulatory dorsomedial PFC. Together these findings suggest a shift in the balance between dorsal and ventral striatal control in cannabis dependence. Similar changes have been observed in animal models and may promote the loss of control central to addictive behavior.

## 1. INTRODUCTION

Converging evidence from animal and human research indicates that maladaptations in fronto-limbic-striatal circuitries drive the loss of control that characterizes addiction (Fineberg et al., 2010; Morein-Zamir and Robbins, 2015). The striatum lies at the core of this circuitry and critically contributes to both, acute drug reinforcement and the transition from voluntary to addictive use that is accompanied by a loss of control (Brand et al., 2016; Everitt and Robbins, 2016; Fineberg et al., 2010).

The striatum contributes to several domains that undergo critical adaptations during the transition to addiction, including associative learning, behavioral control, incentive salience and reward (Haber, 2016; Robbins et al., 2012). The functional heterogeneity of the striatum and its multifaceted contributions to addiction are mirrored in its complex organization into distinct subregions that communicate with the entire cortex via subregion-specific fronto-striatal loops (Haber, 2016). Based on their specific functions and projections, the striatum is divided into the ventral striatum (VS), which includes reward processing hubs such as the nucleus accumbens, and the dorsal striatum (DS) that exhibits strong connections with dorsolateral and dorsomedial prefrontal regions engaged in regulatory control (Di Martino et al., 2008; Haber, 2016; Postuma and Dagher, 2006). This seggregation has important implications for the transition to addiction, with influential conceptualizations proposing that the loss of control is accompanied by a shift between ventral and dorsal striatal control of behavior (Everitt and Robbins, 2016; Jentsch and Taylor, 1999).

Partly driven by the ongoing discussion about cannabis legalization and concomitantly increasing demand for treatment of cannabis-dependence (EMCDDA, 2008; Volkow et al., 2016) a number of studies have examined effects of cannabis use on the brain (overview see e.g. Lorenzetti et al., 2016; Weinstein et al., 2016). In line with the functions of the striato-frontal circuitry and its proposed contribution to addiction task-based neuroimaging studies demonstrated exaggerated striatal reactivity to cannabis cues (Filbey et al., 2016), reward anticipation (Jager et al., 2013) and decreased frontal activity during executive control (Weinstein et al., 2016; Wrege et al., 2014; Yanes et al., 2018) in chronic users. This pattern mirrors mal-adaptations that have been observed across addictive disorders (Brand et al., 2016; Goldstein and Volkow, 2011).

Despite the important contributions of these studies to characterize mechanisms of cannabis dependence, the task-based approach does not allow direct evaluation of the ventral to dorsal striatal shift due to (1) limitation of task-based fMRI to regions engaged by the experimental paradigm (Weinstein et al., 2016), and (2) stimulus- and context-dependence of striatal alterations in cannabis dependence (Gilman, 2017; Zimmermann et al., 2017b). Resting state fMRI functional connectivity (rsfMRI-FC) approaches allow a more holistic assessment of functional alterations in the absence of task or contextual modulation. Moreover, these approaches have a high sensitivity for transient and lasting striatal alterations on the subregion-specific network level (Di Martino et al., 2011; Di Martino et al., 2008). Two recent studies applied rsfMRI-FC to determine lasting distruption in the fronto-striatal functional circuitries in cannabis dependence (Blanco-Hinojo et al., 2017; Zimmermann et al., 2017b). However, findings were limited by the confirmatory nature of the rsfMRI-FC analysis (Zimmermann et al., 2017b) or the use of literature-based striatal seed regions (Blanco-Hinojo et al., 2017) that have a high sensitivity to capture sub-region specific functional networks in healthy subjects (Di Martino et al., 2008) but might not be sensitive to the specific alterations related to cannabis dependence. Moreover, the ventral to dorsal shift has not been explicitly examined in these studies.

The present study therefore employed a two step approach to (1) precisely map striatal alterations in cannabis-dependent individuals using network classification and, (2) to determine whether a ventral to dorsal striatal shift in connectivity with reward and and regulatory control regions can be observed. In an initial step intrinsic striatal alterations were mapped using a novel fully data-driven network classification approach that operates independently of a priori assumptions about striatal architecture at the voxel level and thus enables higher sensitivity to detect striatal functional changes (Martuzzi et al., 2011; Rubinov and Sporns, 2010). Next, altered functional communication of the determined striatal subregions was examined on the whole-brain network level (for a similar approach see Walpola et al., 2017). To this rsfMRI data was acquired in n = 28 participants that fulfilled the DSM-IV criteria for cannabis dependence and n = 28 matched healthy controls. To control for subacute effects of cannabinoid metabolites (Vandevenne et al., 2000), craving (Copersino et al., 2006) and rapid neural recovery of homeostatic receptor adaptations after cessation of chronic cannabis use (Hirvonen et al., 2012) participants underwent 28-days of cannabis abstinence before acquisition of the MRI data. Comparable abstinence periods have been employed in previous studies to differentiate enduring (long-term) effects of cannabis use from transient (sub-acute) alterations and rapid neural recovery (overview in Crean et al., 2011; Ganzer et al., 2016; Schreiner and Dunn, 2012).

Based on current neurobiological frameworks suggesting that the transition to addiction is accompanied by regional-specific neuroplastic changes in the ventral and dorsal striatum and accumulating evidence from task-based neuroimaging studies suggesting that chronic cannabis use is associated with abberant reward and regulatory control processes (Everitt and Robbins, 2016; Weinstein et al., 2016; Wrege et al., 2014; Yanes et al., 2018) we expected to observe corresponding maladaptations on the level of the network level organization of the brain. Specifically we hypothesized that (1) global connectivity differences between the groups specifically map to the dorsal and ventral striatum, reflecting regional-specific striatal adaptations, and (2) a shift in the network communication of the striatal subregions reflecting increased communication within fronto-striatal pathways related to reward processing and concomitantly decreased communication in pathways related to cognitive control.

## 2. MATERIALS and METHODS

### 2.1 Participants

Participants in the present study partly overlapped with the data presented in (Zimmermann et al., 2017b). To enhance the statistical power sample size was increased to n = 28 male cannabis-dependent subjects and n = 28 non-using controls. To control for potential confounding influences of menstrual cycle on striatal resting state activity as well as complex interactions between menstrual cycle and addiction-related alterations in fronto-striatal functioning (Franklin et al., 2015; Wetherill et al., 2016; Wiers et al., 2016), only male subjects were included (for a similar approach see Zimmermann et al., 2017a).

All cannabis users were diagnosed with a current cannabis dependence according to DSM IV during the 18 months before the examination. To facilitate the determination of long-term alterations associated with cannabis dependence that persist beyond subacute cannabinoid metabolites (Vandevenne et al., 2000), craving (Copersino et al., 2006) and rapid neural recovery (Hirvonen et al., 2012) participants were required to abstain from cannabis for 28 days prior to the assessment.

At the time point of recruitment, most cannabis-dependent subjects were active cannabis users or in early phases of abstinence. Inclusion criteria for all participants were: (1) age between 18 and 35 years, (2) right-handedness, and (3) negative urine toxicology for cannabis and other illicit drugs (immunoassay, substance/cut-off per ml: THC/50ng, amphetamines/500ng, cocaine/300ng, methamphetamine/500ng, MDMA/300ng, opiates/300ng, methadone/300ng). Exclusion criteria for all participants were: (1) profound DSM-IV axis I or axis II disorders, such as psychotic or bipolar symptoms, (2) Beck Depression Inventory (BDI-II) score > 20 indicating a moderate depression, (3) medical disorder, (4) current/regular medication, and (5) use of other illicit substances > 75 lifetime occasions. One user reported having used cannabis on one occasion 14 days before the experiment, but was included due to a negative urine toxicology. Controls were included if their cumulative lifetime use of cannabis was < 15g.

Anxiety, mood and concentration were assessed on the day of the examination using validated scales including the social interaction anxiety scale (SIAS) (Mattick and Clarke, 1998), the state-trait-anxiety inventory (STAI) (Spielberger, 1989), positive and negative affect schedule (PANAS) (Crawford and Henry, 2004) and the d2 test of attention (Brickenkamp and Zillmer, 1998). These measures allowed control for potential confounding effects of common withdrawal symptoms. Differences in potential confounders were examined by independent sample t-test; in case of a non-normal distribution of the underlying data non-parametric analyses were used.

Cannabis-dependent subjects were recruited in cooperation with the LVR Clinics Bonn, Germany, Addiction Department that offers a specialized treatment program for cannabis dependence. Written informed consent was obtained and study protocols were approved by the local ethics committee (Medical Faculty, University of Bonn), adhered to the latest revision of the Declaration of Helsinki and were pre-registered at clinicaltrials.gov (NCT02711371).

Three cannabis users were excluded due to a history of exceeding co-use of other illicit drugs and one was excluded due to excessive head movement during fMRI, leading to a final sample of n = 24 cannabis users and n = 28 healthy controls.

### 2.2 MRI Data acquisition

Functional and structural MRI was acquired using a Siemens TRIO 3-Tesla system with a 32-channel head coil. Resting-state fMRI data was acquired before any tasks using a T2*-weighted echo-planar imaging (EPI) pulse sequence (repetition time = 2580ms, echo time = 30ms, number of slices = 47, slice thickness = 3.5mm, no gap, field of view = 224×224mm2, resolution = 64x64, flip angle = 80°, number of volumes = 180). To improve spatial normalization and exclude participants with apparent brain pathologies a high-resolution whole-brain volume T1-weighted images was acquired in addition using a 3D spoiled gradient echo pulse sequence (repetition time = 1660ms, echo time = 2.54ms, flip angle = 9°, field of view = 256×256mm2, acquisition matrix = 256×256, thickness = 0.8mm, number of slices = 208).

Participants were instructed to lie still and relax with their eyes closed while thinking of nothing in particular, yet not to fall asleep during the resting state acquisition.

### 2.3 Preprocessing of the functional resting state data

Resting state fMRI data was preprocessed using standard SPM, AFNI and FSL routines in combination with advanced independent component analysis (ICA-AROMA, Pruim et al., 2015b). The first five volumes were discarded to allow MRI T1 equilibration. The remaining volumes were slice-time corrected, spatially realigned to the first volume, and unwarped to correct for nonlinear distortions possibly related to head motion or magnetic field inhomogeneity using SPM12 (http://www.fil.ion.ucl.ac.uk/spm/software/spm12/). The functional time-series were further processed using the FMRIB Software Library (FSL, http://www.fmrib.ox.ac.uk/fsl), including non-brain removal using BET (Smith, 2002), spatial smoothing with a Gaussian kernel of 6 mm full width at half maximum, and grand-mean scaling. Subsequently, the data was submitted to an independent component analysis for automatic removal of motion artifacts (ICA-AROMA, Pruim et al., 2015b)a procedure that has been shown to minimize the impact of motion on functional connectivity metrics and decrease the loss in temporal degrees of freedom compared to spike regression and scrubbing (Pruim et al., 2015a). Next, mean signals from white matter (WM) and cerebrospinal fluid (CSF), linear and quadratic drift across time, together with a series of sine and cosine functions removing all frequencies outside the range (0.01–0.08 Hz), were regressed out in a single regression step using AFNI’s 3dTproject. Similar to the preprocessing protocol used by Pruim et al. (2015b), WM and CSF time-series were derived by determining the mean time-series over voxels within predefined subject-specific WM and CSF masks. To obtain these masks, we first thresholded the MNI152 average CSF and WM priors maps (95% of the robust range) and subsequently registered to native EPI space.

Likewise, we applied FSL FAST (Zhang et al., 2001) to the individual T1 images to derive a CSF and WM probability map and thresholded at 95% followed by registration to native EPI space. Multiplication of both masks resulted in the respective conservative CSF and WM masks. Finally, functional MRI data were registered to T1 and standard MNI152 space using Boundary-Based Registration as implemented in FLIRT (Jenkinson et al., 2002; Jenkinson and Smith, 2001) and FNIRT nonlinear registration (Andersson et al., 2007) with 6 and 12 df, respectively and interpolated to 2 × 2 × 2mm^3^.

### 2.4 Quality control for motion artifacts

One participant (cannabis user) was excluded due to excessive head movement (> 2.5 mm or 2.5° absolute motion over the whole scan). Mean-frame-wise displacement (FD, Power et al., 2012) and spike control was employed for data quality assurance and motion control. FD was calculated according to with the raw functional data for the remaining participants.

Importantly, the two groups (controls and cannabis users) did not differ in mean FD (controls: Mean ± SD = 0.087 ± 0.032, range from 0.044 to 0.145; cannabis users: Mean ± SD = 0.083 ± 0.024, range from 0.042 to 0.142, *p* = 0.568, Cohen’s d = 0.160). We also defined a spike in head movements as FD > 0.3 mm. Less than 7% scans were regarded as spikes in all of the participants and the two groups did not differ in the number of spikes (control: Mean ± SD = 2.427 ± 3.501, range from 0 to 11; user: Mean ± SD = 1.750 ± 2.723, range from 0 to 11. *p* = 0.445, Cohen’s d = 0.214).

### 2.5 Determining altered striatal connectivity in cannabis dependence using the Intrinsic Connectivity Contrast (ICC)

To promote an unbiased determination of striatal subregions exhibiting altered functional connectivity related to cannabis dependence, a data-driven network level approach was implemented (Walpola et al., 2017). To this end, whole-brain voxel-to-voxel connectivity was computed for each striatal voxel using the intrinsic connectivity contrast (ICC), an index similar to “degree” in graph theory but without the need for a correlation threshold, which reflected the average r^2^ of a given voxel with all other brain voxels (Martuzzi et al., 2011; Rubinov and Sporns, 2010). The corresponding ICC analysis was implemented by weighting the connections of a given voxel with every other voxel in the brain (restricted to the SPM gray matter mask > 0.3) by their r^2^ value (Martuzzi et al., 2011), thus both negative and positive correlations will be captured in positive ICC values.

Next, resulting ICC maps were used for discriminating between cannabis-dependent subjects and healthy controls using a multivoxel pattern analysis (MVPA) with a linear kernel support vector classifier as implemented in LIBSVM (www.csie.ntu.edu.tw/~cjlin/libsvm). A searchlight procedure with a three-voxel radius was used to provide measures of classification accuracy in the neighborhood of each voxel in the striatum mask which was obtained from the Harvard-Oxford Subcortical Structural Atlases (subcortical HO atlas, Harvard Center for Morphometric Analysis) by combining the accumbens, caudate, putamen and pallidum masks (thresholded at 0% population probability) that were further masked by the gray matter mask to remove voxels with low gray matter signal. Classification was evaluated by a ten-fold cross-validation during which all participants were randomly assigned to 10 -subsamples of 5 or 6 participants using MATLAB’s cvpartition function. In each iteration, the optimal hyperplane was computed based on the multivariate pattern of ICC values of 47 or 46 participants and evaluated by the excluded 5 or 6 participants. The default cost parameter (C = 1) was automatically corrected for unbalanced groups by adjusting weights inversely proportional to group frequencies in the training data. The iteration was repeated 10 times with each group being the testing set once and then the average classification accuracy of each sphere was assigned to the center voxel in the sphere. The training set was scaled to [-1, 1], and the testing set was scaled using the same scaling factors before applying SVM (Hsu et al., 2003). To avoid a potential bias of train-test splits, the cross-validation procedure was repeated 5 times by producing different splits in each repetition and the resultant maps were averaged to produce a convergent estimation. To test whether the resultant measures exceeded chance level, we used permutation tests to simulate the probability distribution of the classification. Briefly, we randomly shuffled the group labels and re-computed these measures (repeated 5,000 times) to build empirical distributions. The resultant maps were then converted to *p* values and family-wise error (FWE) corrected for multiple comparisons.

### 2.6 Follow-up seed-to-voxel functional connectivity

Given that ICC is predominantly an exploratory strategy that allows a data-driven precise mapping of striatal regions characterized by their functional connectivity profile differences between the groups functional connectivity analysis was used to determine the associated networks that contribute to these differences (for a similar approach see Martuzzi et al., 2011; Walpola et al., 2017). Importantly, the rationale for the present analyses was based on addiction animal models that indicate a ventral-to-dorsal shift in striatal processing underlying the development of addiction (Everitt and Robbins, 2016). In line with our expectations, the MVPA on the ICC revealed group differences in the global connectivity profiles of ventral and dorsal striatal regions (details see results section). Consequently, the major aim of the follow-up functional connectivity analysis was to determine regions that shift communication with ventral vs dorsal striatal nodes in drug dependence. To this end, two 6-mm-radius spheres (masked by the striatum mask and the gray matter mask) centered at the peak voxels derived from the MVPA results (dorsal, ventral striatum) were used as seed regions, and Fisher Z-transformed image maps were then created by calculating the correlation coefficient of each voxel in the brain with the mean time-series of the seed regions. To determine the shift in ventral vs dorsal striatal networks associated with dependence, the subject-level seed-specific Z-maps were submitted to a 2-way mixed ANOVA with seed (VS and DS) entered as a within-subject factor and group (controls and cannabis users) entered as a between-subject factor employed in the FSL’s Randomise Tool. Corrections for multiple comparisons were conducted using permutation-based inferences (10,000 permutations) with Threshold-Free Clustering Enhancement (TFCE) which provides a strict control while improving replicability (Chen et al., 2017; Smith and Nichols, 2009). For regions showing significant interaction effects (F-test), the mean Fisher z-transformed correlation coefficients were extracted from the underlying anatomical regions for further post hoc analysis within SPSS.

### 2.7 Validation using literature-based ventral and dorsal striatal seeds

In addition, to validate the seed-to-voxel functional connectivity results, functional connectivity analyses were then rerun using predefined VS and DS masks. An anatomical VS mask was defined as nucleus accumbens (subcortical HO atlas) with the probability threshold set to 30% (228 voxels in total). Following Tinnermann et al. (2017), coordinates from different studies (Daw et al., 2006; Fitzgerald et al., 2011; Luijten et al., 2017; Schönberg et al., 2007; Sweitzer et al., 2016) were averaged to create the mean coordinates of reported DS peak voxels. For those lateralized coordinates we multiplied x coordinates by -1 to cover both hemispheres. The DS mask was then created by drawing two 6-mm-radius spheres centered at ± 13, 11, 14 (for bilateral DS) and masked by the striatum mask (223 voxels in total). To control for non-gray matter signal, including WM and CSF, both VS and DS masks were masked by the gray matter mask.

### 2.8 Functional characterization of the determined networks

To support the functional characterization of the networks that exhibit a dependence-associated shift in VS versus DS communication, the NeuroSynth decoder function (http://neurosynth.org/decode/) was used to employ a large-scale automated meta-analysis map. The top 25 terms (excluding terms for brain regions) ranked by the correlation strengths between the regions exhibiting dependence-associated alterations and the meta-analytic map were visualized using word cloud with the size of the font scaled by correlation strength.

## 3. RESULTS

### 3.1 Group Characteristics

The groups were comparable with respect to important potential confounders including socio-demographics, attention and the use of licit drugs (**Table 1**). The lack of significant group differences in attention, mood and state anxiety (**Table 1**) argues against strong confounding effects of cannabis withdrawal. As expected cannabis users reported greater lifetime experiences with other illicit drugs (**Table 1**) than controls. Cannabis use parameters are reported in **Table 2**. In post-MRI interviews none of the subjects reported having fallen asleep during the scan.

**Table 1.** Group characteristics and drug use parameters. aMann-Whitney-U test, *Median(Range), ** Prescription medicinal use

| Measure | Marijuana Users<br>M (SD) | Controls<br>M (SD) | p |
| --- | --- | --- | --- |
| Age (years) | 24.00(3.46) | 23.39(2.86) | 0.49 |
| Education (years) | 14.50(11.00-22.00)* | 14.25(12.00-19.00)* | 0.75<sup>a</sup> |
| d2 concentration performance | 196.46(39.61) | 205.43(50.57) | 0.49 |
| STAI-state | 34.17(7.70) | 31.07(7.71) | 0.16 |
| STAI-trait | 35.87(8.40) | 32.64(7.22) | 0.14 |
| SIAS | 16.50(8.00-46.00)* | 16.00(5.00-33.00)* | 0.24<sup>a</sup> |
| Number of smokers | N = 22 | N = 23 |  |
| Years of nicotine use | 8.23(4.49) | 6.24(3.72) | 0.11 |
| Cigarettes per day | 8.00(2-20.00)* | 8.00(1-30.00)* | 0.96<sup>a</sup> |
| Pack-year | 4.16(3.59) | 3.42(3.84) | 0.50 |
| Alcohol occasions per week | 1.00(0-4.00)* | 1.00(0-4.00)* | 0.74<sup>a</sup> |
| Alcohol units per week | 5.00(0-24.00)* | 4.90(0-18.00)* | 0.92<sup>a</sup> |
| PANAS-positive | 33.00(16.00-42.00)* | 32.50(22.00-48.00) | 0.67<sup>a</sup> |
| PANAS-negative | 11.00(10.00-25.00)* | 10.00(10.00-14.00)* | 0.10<sup>a</sup> |
| Past ecstasy use | N = 15 | N = 2 |  |
| Lifetime occasions ecstasy | 10.00(1-75)* | (1, 8) | - |
| Past cocaine use | N = 11 | N = 0 |  |
| Lifetime occasions cocaine | 6.00(1-70)* | - | - |
| Past amphetamine use | N = 15 | N = 1 | - |
| Lifetime occasions amphetamine | 20.00(1-75)* | 30.00 | - |
| Past hallucinogen use | N = 9 | N = 0 | - |
| Lifetime amount hallucinogen | 6.00(1-50)* | - | - |
| Past opiate use | N = 3 | N = 1 | - |
| Lifetime occasions opiate | (1, 1, 4) | 30.00** | - |
| Past marijuana use | N = 24 | N = 22 | - |
| % Lifetime marijuana dependence | 100% | 0% | - |

**Table 2.**
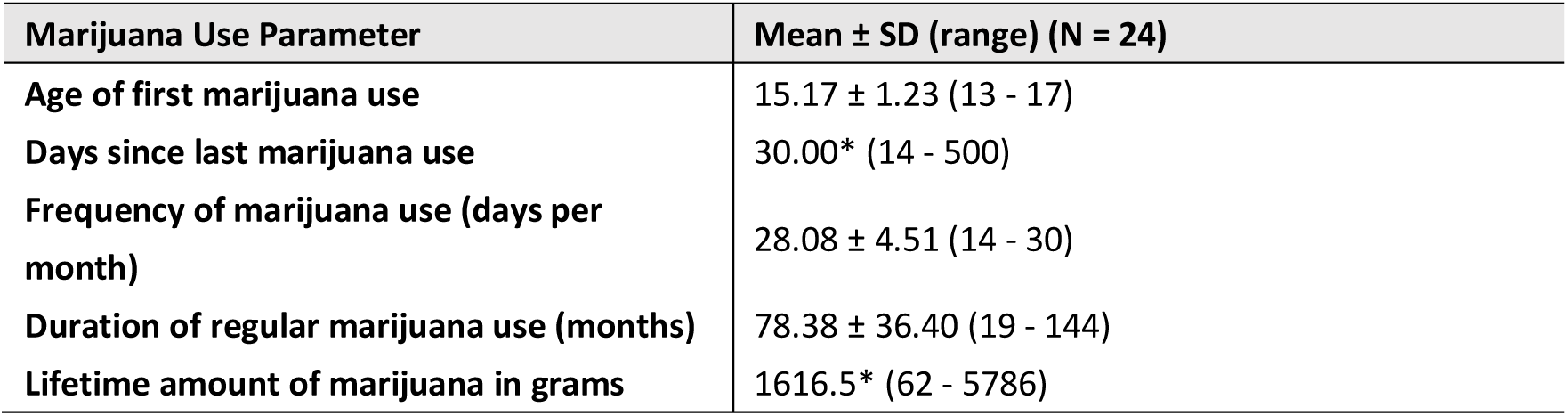
Marijuana use parameters. *Median

### 3.2 Intrinsic connectivity contrast

The searchlight MVPA identified brain regions within the striatum that exhibit altered global connectivity patterns in cannabis users compared to controls. Specifically the groups could be reliably discriminated in two regions that mapped onto the VS (peak voxel coordinates, 10, 12, -10 mapping onto the right nucleus accumbens, accuracy = 84.62%, voxel-wise FWE-corrected *p* = 0.04) and the DS (peak voxel coordinates, -12, 14, 16 mapping onto the left caudate nucleus, accuracy = 85.38%, voxel-wise FWE-corrected *p* = 0.026) (**Figure 1**).

**Figure 1.**
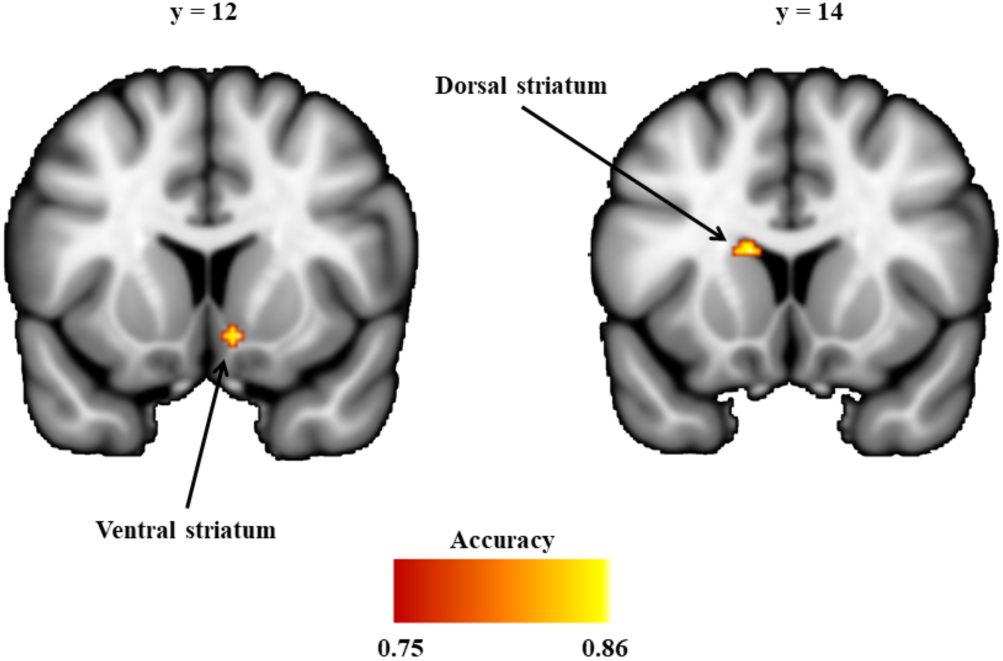
Striatum regions discriminating cannabis dependent subjects and non-using controls. Images are thresholded at accuracy >= 0.75 (approximate p < 0.001, uncorrected) and cluster size > 5 for display purpose.

### 3.3 Follow-up analysesseed-to-voxel functional connectivity

The main effects of seed reflecting the differential connectivity patterns of the VS vs DS seeds further confirmed the mapping of the MVPA to the VS and DS: (1) the VS seed showed stronger connectivity with ventral regions of the frontal cortex including rostal ACC and orbitofrontal regions, whereas the DS seed exhibited stronger connectivity with the dorsal frontal regions including dorsolateral prefrontal cortex, a pattern resembling previous human and animal research (Di Martino et al., 2008; Haber, 2016). (2) There was a strong overlap between the networks identified for the VS and DS regions determined in the present study and the literature-based seeds from the validation analysis (**Figure 2**).

**Figure 2.**
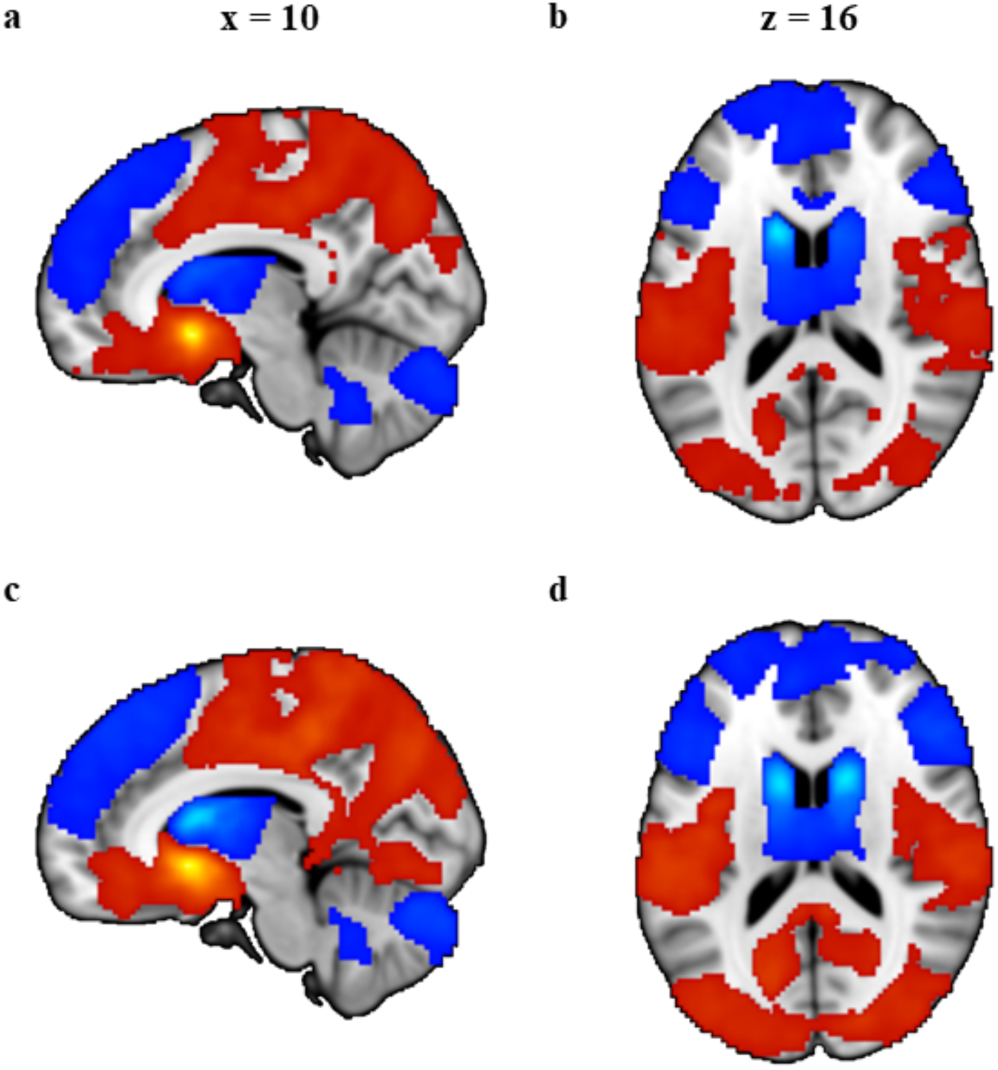
Brain regions showing significant main effect of seed on functional connectivity. (a) and (b) are results from data-driven seeds and (c) and (d) are results from validation seeds. Red color indicated stronger functional connectivity with VS than DS and blue color indicated stronger functional connectivity with DS than VS. All images were thresholded at *p* < 0.05, TFCE corrected.

Examining differences in seed-based functional connectivity between the groups revealed no regions showing a significant main effect of group, arguing against unspecific striatal network changes in cannabis dependence. However, significant group x seed interaction effects were found mirroring a shift in network level communication of the striatal subregions in cannabis dependence. The interaction effects were observed in connectivity with the dorsomedial prefrontal cortex (dmPFC) and rostral anterior cingulate cortex (rACC) across both analyses (**Table 3 and Figure 3**).

**Table 3.**
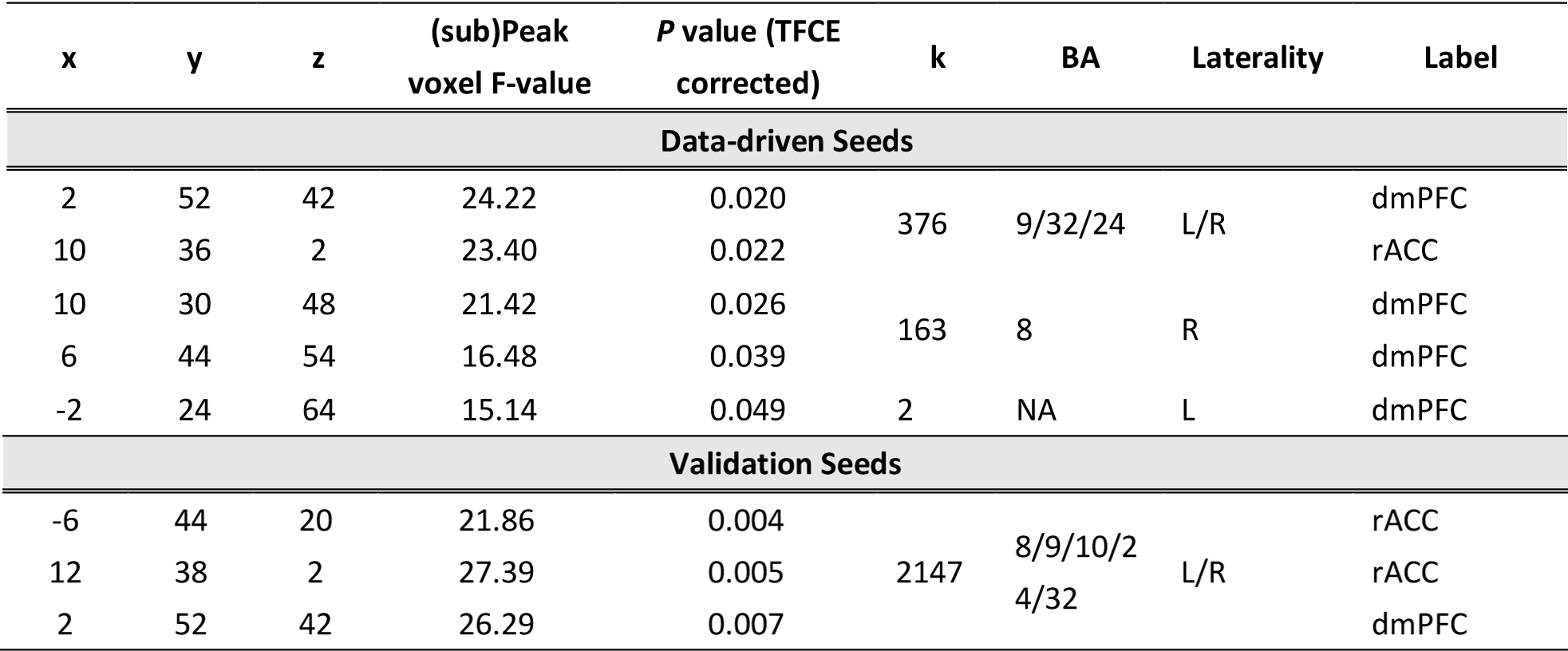
Brain regions showing significant group (control vs user) by seed (ventral striatum vs dorsal striatum) interaction. x, y, z: MNI-coordinates; k: cluster size; BA: brodmann area; dmPFC: dorso-medial prefrontal cortex; rACC: rostral anterior cingulate cortex.

**Figure 3.**
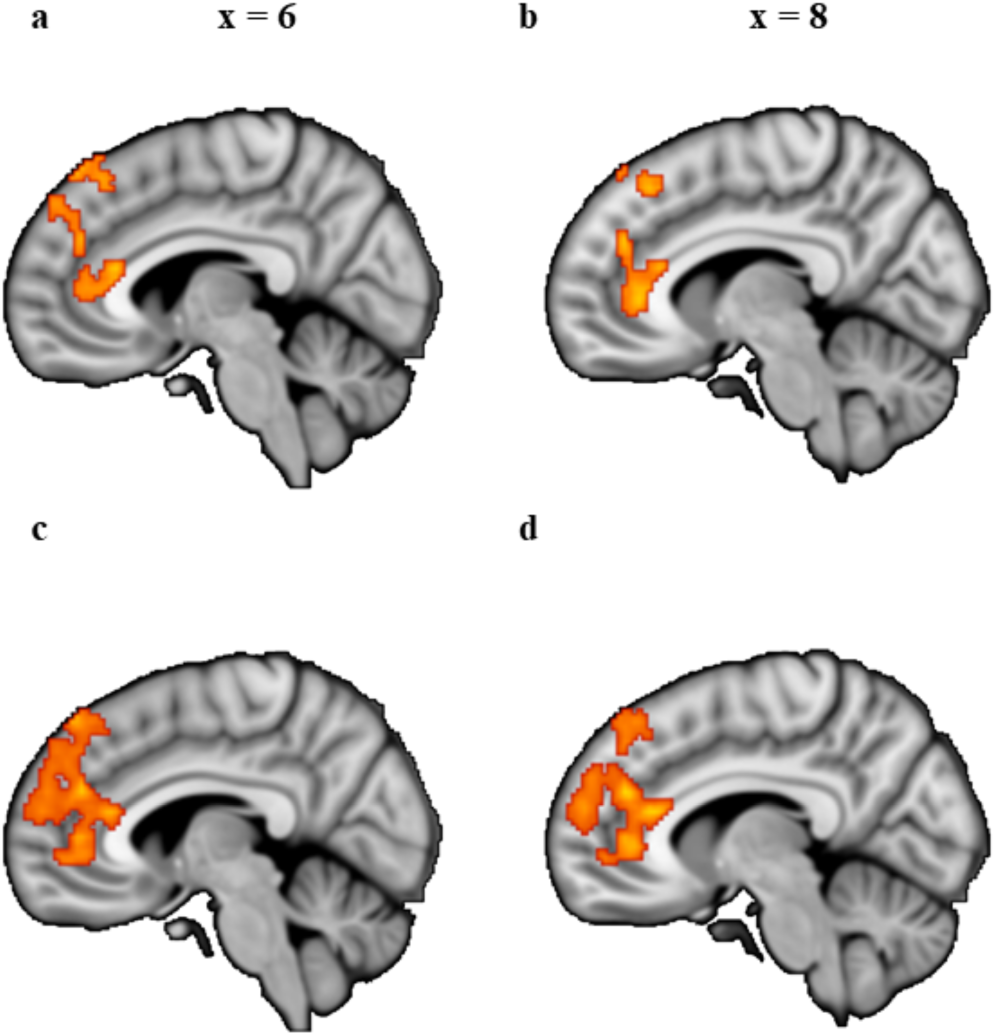
Brain regions showing significant group (control vs user) by seed (ventral vs dorsal striatum) functional connectivity interaction. (**a**) and (**b**) are results from data-driven seeds and (**c**) and (**d**) are results from validation seeds. All images are thresholded at *p* < 0.05, TFCE corrected

### 3.4 Post hoc analyses

To further disentangle the direction of the changes using post hoc comparisons, mean Fisher z-transformed correlation coefficients were extracted from anatomical masks referring to the dmPFC and rACC. Masks of the respective regions were defined using structurally defined regions. To obtain the rACC mask, we first thresholded the anterior cingulate gyrus (HO atlas) at 30% to remove voxels with low probability and the posterior boundary was then delineated as one slice anterior to the coronal plane where the connection of the corpus callosum in each hemisphere was no longer connected (Asami et al., 2008; McCormick et al., 2006). Similar, to obtain the DMPFC mask, we thresholded the superior frontal gyrus (HO atlas) at 30% probability, excluding voxels further than 10 mm from the midline of the brain (de la Vega et al., 2016) or posterior to the anterior boundary of supplementary motor area. As displayed in **Figure 4a and 4c**, post hoc pairwise comparisons revealed attenuated negative correlation (anti-correlation) between dmPFC and VS (*t*_(50)_ = 3.23, *p* = 0.002, Cohen’s d = 0.901), as well as decreased positive connectivity between dmPFC and DS (*t*_(50)_ = -2.33, *p* = 0.024, Cohen’s d = 0.649) in cannabis users relative to controls. Post hoc comparisons on the rACC demonstrated increased positive functional connectivity with the VS (*t*_(50)_ = 2.32, *p* = 0.025, Cohen’s d = 0.644) and concomitantly decreased positive functional connectivity with the DS (*t*_(50)_ = -2.69, *p* = 0.009, Cohen’s d = 0.751) in cannabis users compared to controls.

**Figure 4.**
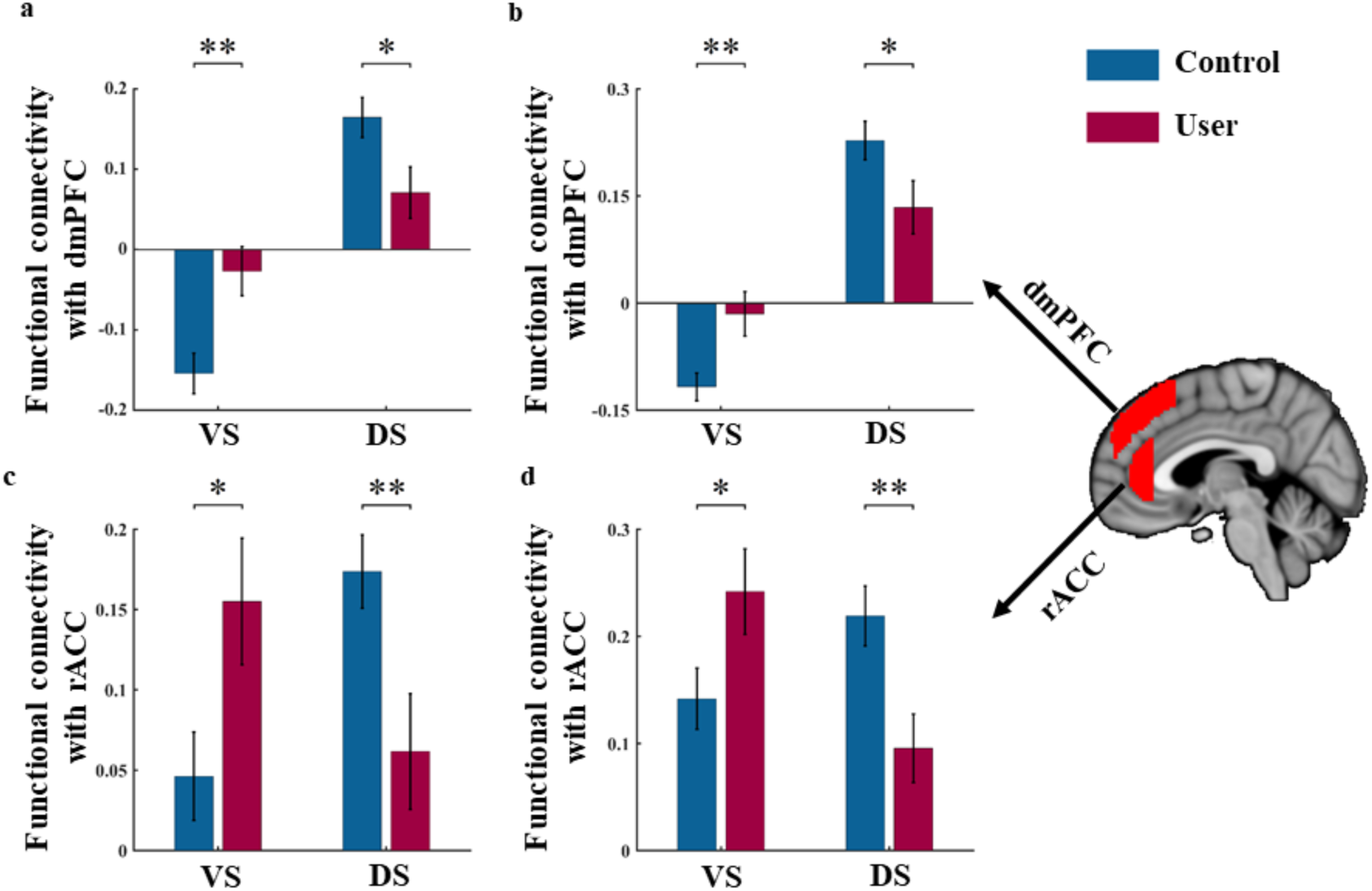
Patterns of functional connectivity associated with a significant group (control vs user) by seed (VS vs DS) interaction. (**a**) and (**c**) are results from data-driven seeds and (**b**) and (**d**) are results from validation seeds. Error bars reflect the SEM. *p < 0.05; **p < 0.01; VS: ventral striatum; DS: dorsal striatum; dmPFC: dorsomedial prefrontal cortex; rACC: rostral anterior cingulate cortex.

### 3.5 Control analyses

Importantly none of the cannabis users fulfilled a dependence for other illicit drugs, and cannabis was their main drug of abuse. However, in line with an increased prevalence of illicit drug use in addicted populations the cannabis group had higher experience with recreational use of other drugs, particularly amphetamine (approx.. 60%). Given that previous studies reported altered striatal functioning in occasional amphetamine users (Schrantee et al., 2016), we additionally compared the altered neural indices between users with a history of amphetamine use versus no history. This additional control analysis did not reveal significant between group differences (all ps > 0.2), arguing against recreational amphetamine use as driver of the observed difference with the control group. Moreover, the control analysis with the literature-derived validation seeds revealed comparable results as the data-driven seeds, further confirming the robustness of the findings (**Figure 4b and 4d**).

### 3.6 Functional characterization of the altered networks

NeuroSynth decoding revealed that terms showing highest correlations with the dmPFC were predominantly referring to control-related or reasoning-related processes (**Figure 5a**), while a predominance of reward-related or default network processes (**Figure 5b**) was found for the rACC.

**Figure 5.**
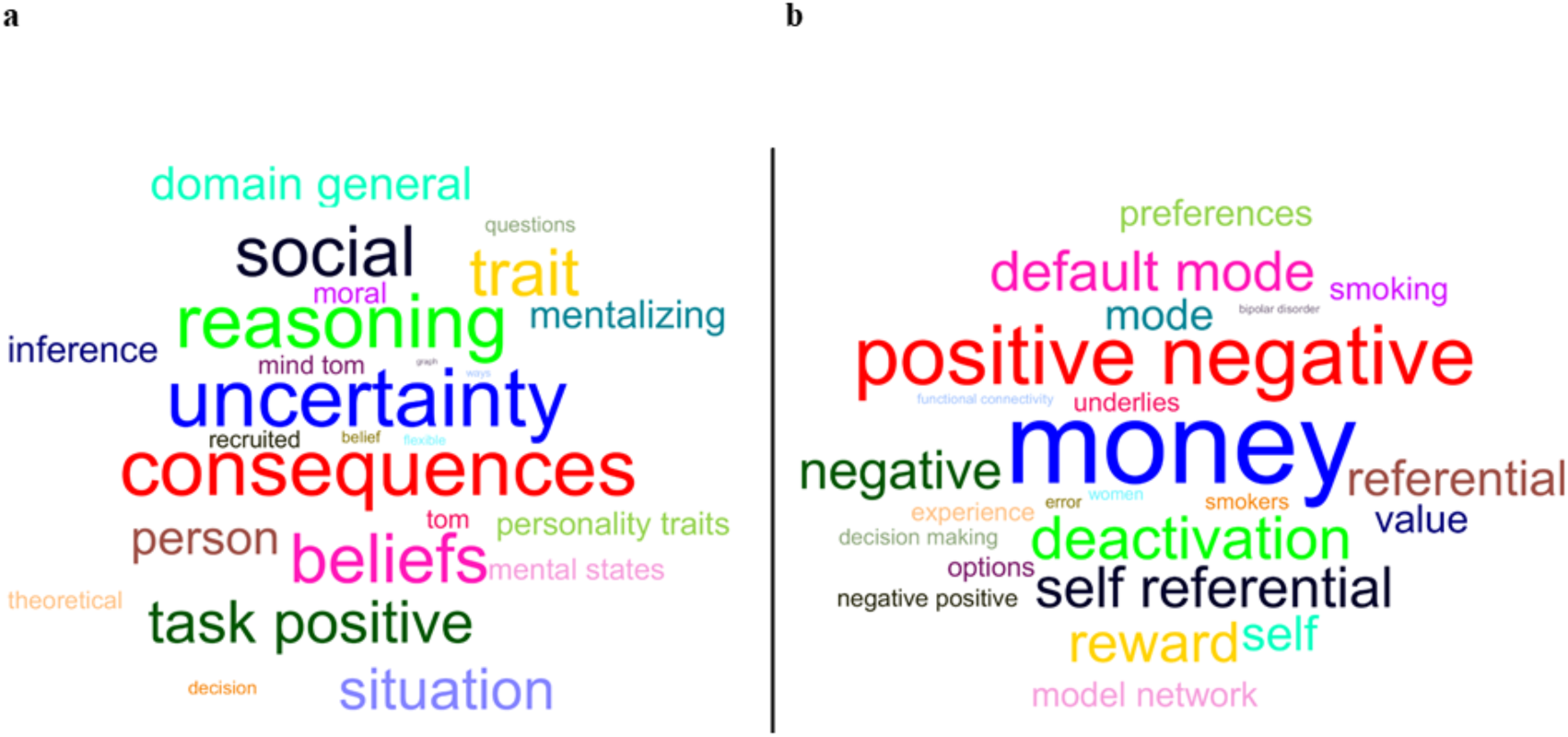
Word cloud showing the correlation strength between the ROIs and the meta-analytic map. **(a)** The top 25 terms associated with dmPFC based on Neurosynth decoding and font size represents relative correlation strength of that term to dmPFC. (**b**) The same information for rACC.

### 3.7 Association between cannabis use parameters and neural alterations

Exploring associations between neural alterations and the age of onset of cannabis use, cumulative exposure and duration of abstinence (days since last use) did not reveal significant associations (all ps > 0.1).

## 4. DISCUSSION

The present study combined an innovative network level approach with resting state fMRI to determine regional-specific striatal alterations and associated changes in functional communication in abstinent cannabis-dependent subjects. In line with animal and human data that emphasize the specific contributions of the VS and DS, subregion-specific differences between cannabis users and controls mapped to the nucleus accumbens and the caudate nucleus. Cannabis users displayed a relative shift between VS and DS communication with fronto-limbic regions that have been consistently involved in reward processing (rACC) and executive / regulatory functions (dmPFC). Aberrant fronto-striatal functional connectivity in the abstinent cannabis users was observed in the absence of a significant main effect in the comparison with the control group. This observation is partly in line with a previous study reporting that altered fronto-striatal connectivity in chronic cannabis users normalizes during 28 days of abstinence (Blanco-Hinojo et al., 2016). However, comparing the shift between the ventral and dorsal striatal functional networks in the present study revealed evidence for a persisting imbalance in the subregion specific fronto-striatal circuits. Together, these findings argue against unspecific long-term striatal changes in cannabis dependence but rather emphasize the importance of both, specific adaptations in dorsal and ventral subregions and a relative shift between the regions.

Moreover, employing a fully-data driven network approach that combines the intrinsic connectivity contrast with a pattern classification demonstrated that the general connectivity patterns of the VS and DS can reliably discriminate between cannabis-dependent individuals and controls. As such, the present data are in line with previous studies reporting altered ventral and dorsal striatal activity in chronic cannabis users during drug cue reactivity, reward processing (Weinstein et al., 2016; Wrege et al., 2014; Yanes et al., 2018) and social decision making (Gilman, 2017). Further, our data emphasize the important contribution of striatal maladaptations in addiction which might reflect a common pathological pathway across addictive disorders. In accordance with a plethora of previous animal and human research suggesting a functional differentiation of the striatum along a dorsal to ventral gradient (Delgado, 2007) and a specific contribution of the striatal subregions to the transition to addiction (Everitt and Robbins, 2016), differences in the intrinsic function of both subregions were demonstrated. Specifically, the nucleus accumbens, a core reward processing node (Delgado, 2007), and the caudate nucleus, strongly engaged in cognitive processes and executive control (Haber, 2016), differed between both groups.

The mapping of the identified regions and their proposed functional contributions was further confirmed by the stronger connectivity of the VS with reward-related limbic and (orbito-)frontal regions, whereas the DS showed a stronger functional interaction with frontal regions such as the dorsolateral prefrontal cortex (dlPFC) that are critically involved in cognitive and regulatory processes. Examining differences in the connectivity profiles of the identified VS and DS regions revealed a relative shift in the connectivity with the dmPFC, a region that has been implicated in regulatory downstream control over the striatum (Kragel et al., 2018; Robbins et al., 2012) and the rACC, a region engaged in reward processing via up-stream signaling of the VS (Haber and Knutson, 2010). As such, the present findings are in accordance with recent quantitative and qualitative reviews suggesting alterations in networks engaged in cognitive control and reward processing in chronic cannabis users (Weinstein et al., 2016; Wrege et al., 2014; Yanes et al., 2018).

More specifically, both striatal regions demonstrated an uncoupling with dmPFC regulatory regions which may reflect deficient inhibitory control which is consistently observed in cannabis users (Wrege et al., 2014). The altered caudate-dmPFC pathway suggests specific deficits in stop-signal inhibition (Robbins et al., 2012), an inhibitory domain that has been shown to be deficient across addictive disorders (Acikalin et al., 2017; Morein-Zamir et al., 2013). By contrast, in cannabis-dependent subjects the rACC exhibited increased connectivity with the VS and concomitantly decreased connectivity with the DS. Increased connectivity in the nucleus accumbens – ACC pathway in response to drug cues has previously been identified as a specific marker for dependent versus non-dependent cannabis users (Filbey and Dunlop, 2014). Activation in the rACC and the adjacent ventromedial PFC have been found to reliably reflect subjective value and reward representations (Acikalin et al., 2017). The pathway with the VS has been associated with the degree of delayed discounting-associated inhibitory control (Li et al., 2013; see also circuit model by Robbins et al., 2012) and deficits in this domain have also been observed in cocaine-dependent individuals (Contreras-Rodríguez et al., 2015). Consistent with the present results on decreased DS-rACC connectivity, a previous study reported attenuated connectivity in this pathway in nicotine-dependent individuals (Sweitzer et al., 2016). This has been interpreted as disengagement of habitual behavior in favor of conscious perception of craving that promotes drug seeking behavior. To this end, altered striatal rACC connectivity may reflect increased sensitivity to rewards as proposed by the incentive salience theory of addiction (Robinson and Berridge, 2000) which has recently been demonstrated in cannabis dependence (Filbey et al., 2016) and an increased sensitivity for internally generated negative feelings related to craving.

The present findings need to be considered in the context of limitations. To facilitate the determination of elementary fronto-striatal alterations in cannabis dependence the study focused on male participants. The rationale was based on an increasing number of studies reporting modulatory effects of menstrual cycle phase on fronto-striatal processing and associated functions (e.g. Diekhof and Ratnayake, 2016; Dreher et al., 2007) as well as recent studies suggesting that addiction-related alterations in these circuits vary as a function of menstrual cycle (Franklin et al., 2015; Wetherill et al., 2016; for cannabis see Wiers et al., 2016). The focus on a male sample allowed to control for these potential confounding variables, however, comes at a cost of a limited generalization of the present findings to cannabis dependent women. In the context of emerging evidence for a higher sensitivity of women to the adverse effects of chronic cannabis use on the brain, including blunted dopaminergic reactivity in the striatum and impaired frontal activity (Paola Castelli et al., 2014; Wiers et al., 2016), future studies need to consider to explicitly examine sex-differences, (2) due to the high prevalence of tobacco co-use in cannabis addiction both samples included a high proportion of tobacco smokers, although the groups were matched with respect to tobacco use parameters complex interactions between cannabis and tobacco use may have contributed to the present findings, and (3) the cannabis sample size was relatively small which limited ability to assess within-group clinical correlations.

Together the present findings suggest a shift in the balance between dorsal and ventral striatal control of behavior in cannabis dependence. Similar changes have been previously validated in comprehensive animal models of addiction and may promote the loss of control central to the transition to addictive behavior.

## FUNDING AND DISCLOSURE

This work was supported by the National Natural Science Foundation of China (NSFC, 91632117; 31530032), the German Research Foundation (DFG, grant: BE5465/2-1, HU1302/4-1), All other authors report no biomedical financial interests or potential conflicts of interest.

